# Dogs’ sensory-motor tuning shapes dog-human vocal interactions

**DOI:** 10.1101/2023.10.18.562860

**Authors:** E. C. Déaux, T. Piette, F. Gaunet, T. Legou, L. Arnal, A-L. Giraud

## Abstract

Within species, vocal and auditory systems co-evolve to converge on a critical temporal acoustic structure that can be best produced and perceived. While dogs cannot produce articulated sounds, they respond to speech, raising the question as to whether this heterospecific receptive ability is shaped by exposure to speech or bounded by their own sensorimotor capacity. Acoustic analyses of vocalisations show that dogs’ main production rhythm is slower than the dominant (syllabic) speech rate, and that human dog-directed speech falls halfway in between. Comparative exploration of neural (electroencephalography) and behavioural responses to speech reveals that comprehension in dogs relies on a slower speech rhythm tracking (delta) than humans’ (theta), even though dogs are equally sensitive to human speech content and prosody. Thus, the dog audio-motor tuning differs from humans’, who vocally adjust their speech rate to this shared temporal channel.

---

Acoustic communication evolves through a dynamic closed-loop phenomenon in which auditory systems are tuned to vocal signals, while in turn vocal production adapts to exploit the capacity of sensory systems ^1–4^. In this fine audio-vocal tuning, temporal acoustic features have a universal ecological relevance, being essential for e.g. vocal recognition ^5,6^ predator avoidance ^7^ or mate choice ^8–10^.

Exploration of the speech system has provided capital insight into the neural bases of this perception/production tuning. Speech rhythms are mechanically constrained by the motor effectors, but also operated within a certain dynamic range to best match perception-action neural rhythms. Thus, the dominant speech rhythm, the syllable rate, is cross-culturally stable ^11^ because it both arises from the interplay of the different articulators ^12,13^ and corresponds to neural theta oscillations, involved in active sensing across species ^14^. In speech perception, the auditory theta rhythm serves to actively interface the acoustics with endogenous neural processes, and the closer the acoustics to this rhythm the more efficient the information transfer. Crucially, the neural theta rhythm can *flexibly* adapt to speech quasiperiodicity via a mechanism referred to as “speech tracking” ^15^, and comprehension critically depends on its precision ^16–21^. Thus speech production and reception tuning has led to a common temporal window of analysis centred on the 4-8Hz range ^22^.

Among animals, dogs, *Canis familiaris*, are one of the most adept at responding to human signals, arguably because of their long co-evolutionary history with us, and are particularly receptive to auditory cues ^23,24^. They can learn upwards of hundreds of words ^25^ and may exhibit fast mapping ^26^ and statistical learning abilities ^27^. Behavioural data consistently show their sensitivity to speech temporal regularities ^28,29^.Yet, dogs lack the vocal/neural system necessary for articulated communication ^30–32^.

This thus begs the question as to how dogs and humans have adapted to each other’s production/perception constraints to produce successful vocal interactions. It may be that the dog neural system has adapted to human speech, or conversely that humans have adjusted their vocal production to exploit the dogs’ neural (auditory) capacity. Interestingly, like parents with infants, dog owners spontaneously use accented speech modulations, referred to as dog-directed speech ^33–35^. Infant-directed speech also accentuates slow rhythm (i.e. delta band) over the syllabic rhythm (i.e. theta band)^36^, which has a positive influence on infant speech tracking ^37^, although a similar analysis of dog-directed speech rhythm is lacking. To address these questions, we first analysed dog vocalisations, as well as adult-(ADS) and dog-directed speech (DDS), to probe whether dogs vocalise at the same or at a different rate than humans, and whether the temporal properties of DDS differ from those of ADS. Second, we compared speech neural processing in dogs and humans using non-invasive electroencephalography (EEG), to investigate whether dogs track speech modulations in sync with their own production system or syllabify speech as humans do. In other words, we explore whether dogs could have evolved a discordance between the capacity of their auditory and vocal systems through phylo- and/or onto-genetic exposure to speech.

## RESULTS

### Natural vocal rate in dogs and humans

Using 143 vocal sequences (30 dogs) including all major vocal classes (barks, growls, howls, snarls and whines ^38^), 106 adult-directed (27 individuals, 10 women) and 149 dog-directed speech sequences (22 individuals, 16 women) spanning five different languages, we found that dogs vocalise at a slower rate than humans (Dogs mean ± SD: 2 ± 1.1 vocalisations/s, ADS: 4 ± 1.9 syllables/s; Tukey-corrected post-hoc pairwise comparison: t=6.8, p<0.001, Figure 1A and 1B). Further, DDS has a slower rate (3 ± 1.6Hz) than ADS (t=3.1, p=0.007), but faster rate than the average dog vocal rate (t=3.9, p=0.006). For a subset of individuals that produced DDS, we found duration-matched ADS sentences, confirming that pet owners slow their speech rate when talking to their dogs (paired t-test: t=2.7, df=11, p=0.02, Cohen’s d= 0.8, Figure 1C). DDS also has higher mean F0 than ADS (t=-2.2, df=11, p=0.05, Cohen’s d= 0.6) confirming previous results ^33,35^. Further analyses of vocal sequences find no significant differences in vocal rate among vocal classes in dogs (F_4,16.1_=1.4, p=0.28) nor among languages in ADS (F_4,19.8_=1.8, p=0.17) or DDS (F_4,9.5_=2, p=0.17, Figure 1E). Thus, the dog’s vocal rate is overall slower than human speech and importantly, pet owners modify not only the spectral but also the temporal feature of their output when speaking to their dogs, in a direction that brings them closer to the natural vocal rate of the latter.

**Figure 1.**
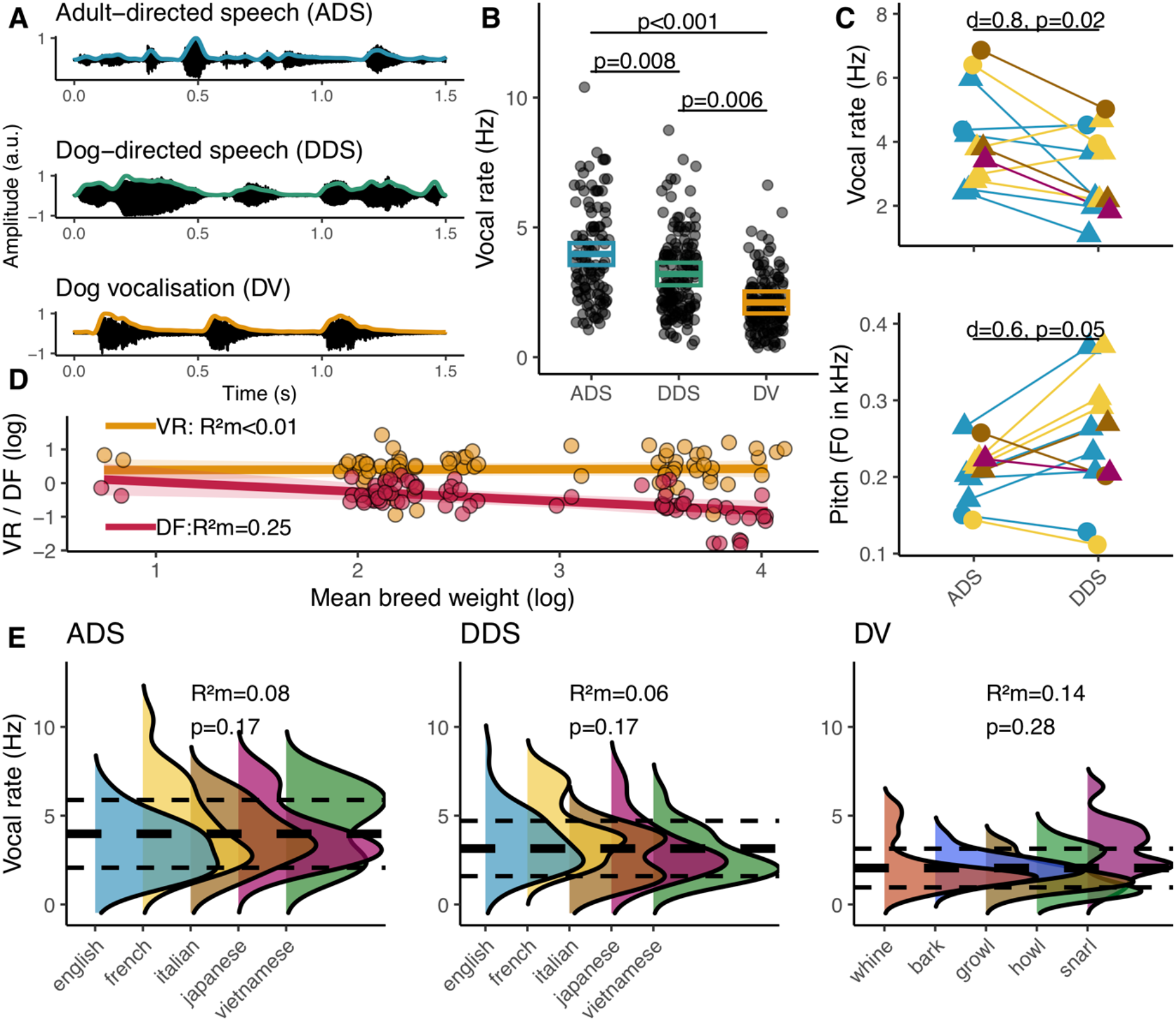
Comparison of dog/human vocal production. A) Oscillograms and, overlaid, envelopes used to compute the vocal rate. B) Model estimates and their 95% CI of vocal rate in dog and human sequences. Black dots are the original observations. C) Vocal rate (Hz) and mean F0 (Hz) for matched ADS and DDS speech sentences. D) Model slope and 95% CI of weight effect on dog vocal rate (VR) and dominant acoustic frequency (DF). E) Density distribution of vocal rate according to vocal classes for dogs and languages for humans. Overall mean (thick dashed line) and SD (thin dashed lines) statistics are displayed.

Furthermore, when exploring other factors known to influence the structure of animal vocal signals ^2^, we found no evidence of large inter-individual differences in vocal rate unlike for the dominant acoustic frequency (table S1) confirming the latter’s functional significance in individual discrimination ^39,40^ and speaking against such selection effects in the former. Concurrently, body weight had no explanatory effect on vocal rate variation (F_1,8.17_=0.03, p=0.88) while it had a strong negative effect on dominant acoustic frequency (F_1,12.07_=6.03, p=0.03), again confirming the known acoustic allometric relationship between the latter two ^41^ and speaking to other types of constraints on vocal rate (Figure 1D)^1^.

### Neural tracking and speech ‘intelligibility’

To investigate auditory neural processes in dogs, we adapted typical human protocols e.g. ^16,18,20^, where speech intelligibility is altered using spectral and temporal modifications of speech stimuli and neural tracking strength is correlated to behavioural measures of intelligibility (Figure 2). Speech streams were composed of words that the dogs had learned to respond to, i.e. command words (e.g. “sit”, “come”) allowing us to use their behavioural responses as an index of ‘intelligibility’ in the sense of a successful stimulus-action relationship. In humans, comprehension was assessed by asking participants to rate word streams on an intelligibility scale. We performed EEG and behavioural experiments on 12 dogs (1-13 years old, seven females) and 12 paired human participants (18-65 years old, six women) with no reported hearing deficits. Four dogs and one human participant were excluded from analyses due to poor EEG signal quality.

**Figure 2.**
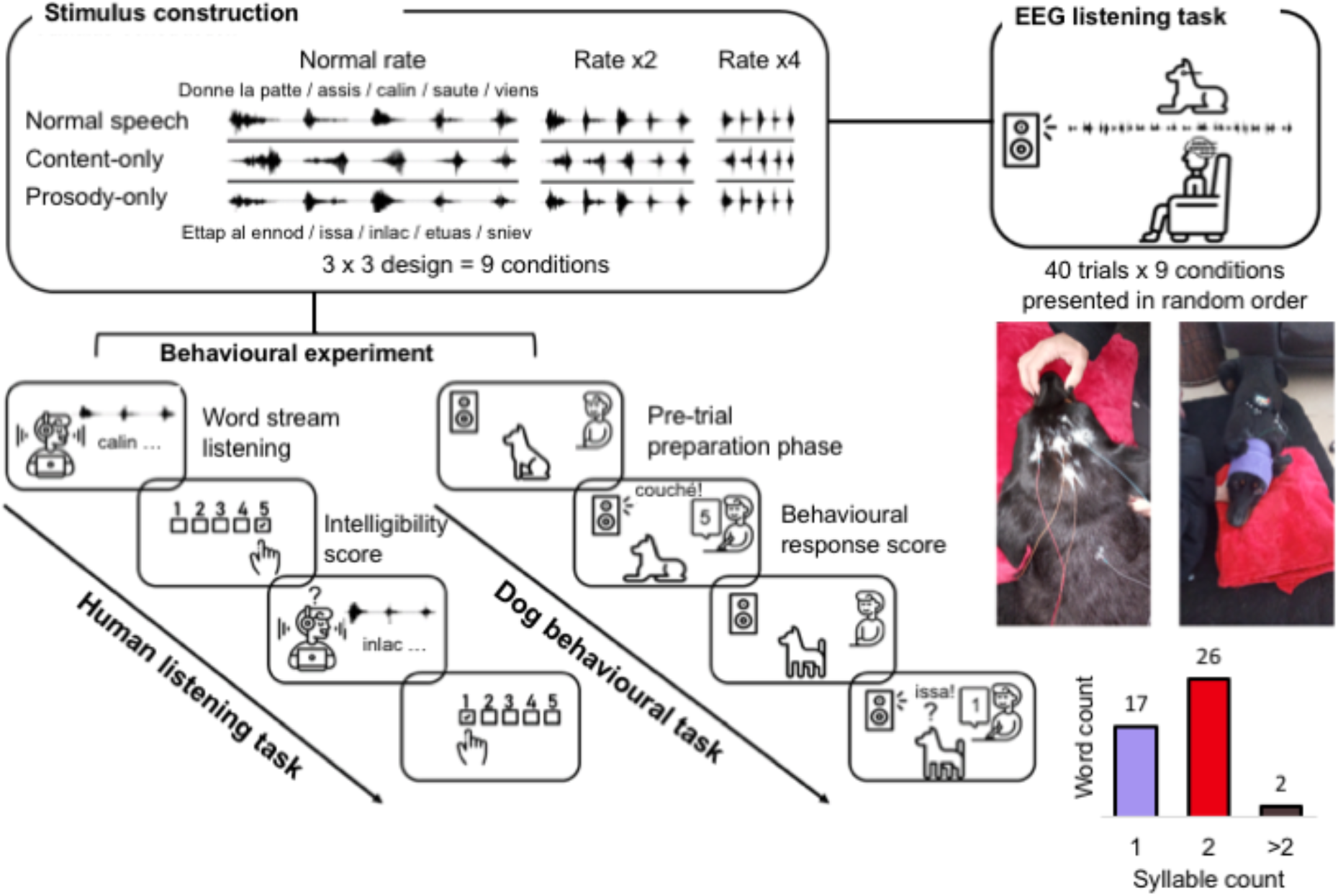
Schematic of the perception study. Word stream stimuli were first constructed by recording dog-specific command words (mostly disyllabic and monosyllabic, cf. small insert) that were appended into a 5-word stream with ∼300 ± 50ms silence intervals. These word streams were altered with regards to 1) speech type: by either removing content (reversed words) or prosodic information (flatten pitch modulation and reversed energy contour) and 2) speech rate: compression by a factor of 2 or 4; amounting to nine word-stream conditions in total. The behavioural experiment consisted of an intelligibility scoring task for humans who listened to the full word stream, and of a playback task for dogs, who heard each word command separately (45 in total) a maximum of 3 times each, while the experimenter and the owner agreed on a behavioural response score. For the EEG experiment dogs were first fitted with 1 to 4 electrodes covered by a headband and linked to an amplifier strapped on their back (photo inserts). They were then instructed to lie down and passively listen to an audio track (broadcasted via a speaker) containing 40 repetitions of each word stream condition. For comparability purposes, human EEG recordings were made under the same experimental conditions.

We first confirmed that modifying speech spectral and temporal features altered both species’ perceptual performances (Figure 3A). Modifying speech rate (main effect: F_2,80_=46.6, p<0.001) and type (main effect: F_2,80_=112, p<0.001) affected speech intelligibility in humans, in an interactive way (rate by type interaction: F_4,80_=10.5, p<0.001). Increasing speech rate decreased speech intelligibility, with a stronger effect in the Normal speech and Content-only conditions. Concurrently, altering speech features had a strong impact on comprehension, particularly when content was removed i.e. the Prosody-only condition. In dogs, speech rate (main effect: F_2,64_=4.9, p=0.01) and type (main effect: F_2,64_=6.4, p=0.003) also impacted speech intelligibility, again interactively (rate by type interaction: F_4,64_=6.9; p<0.001), with intelligibility dropping as speech rate increased, but only in the normal speech type condition.

**Figure 3.**
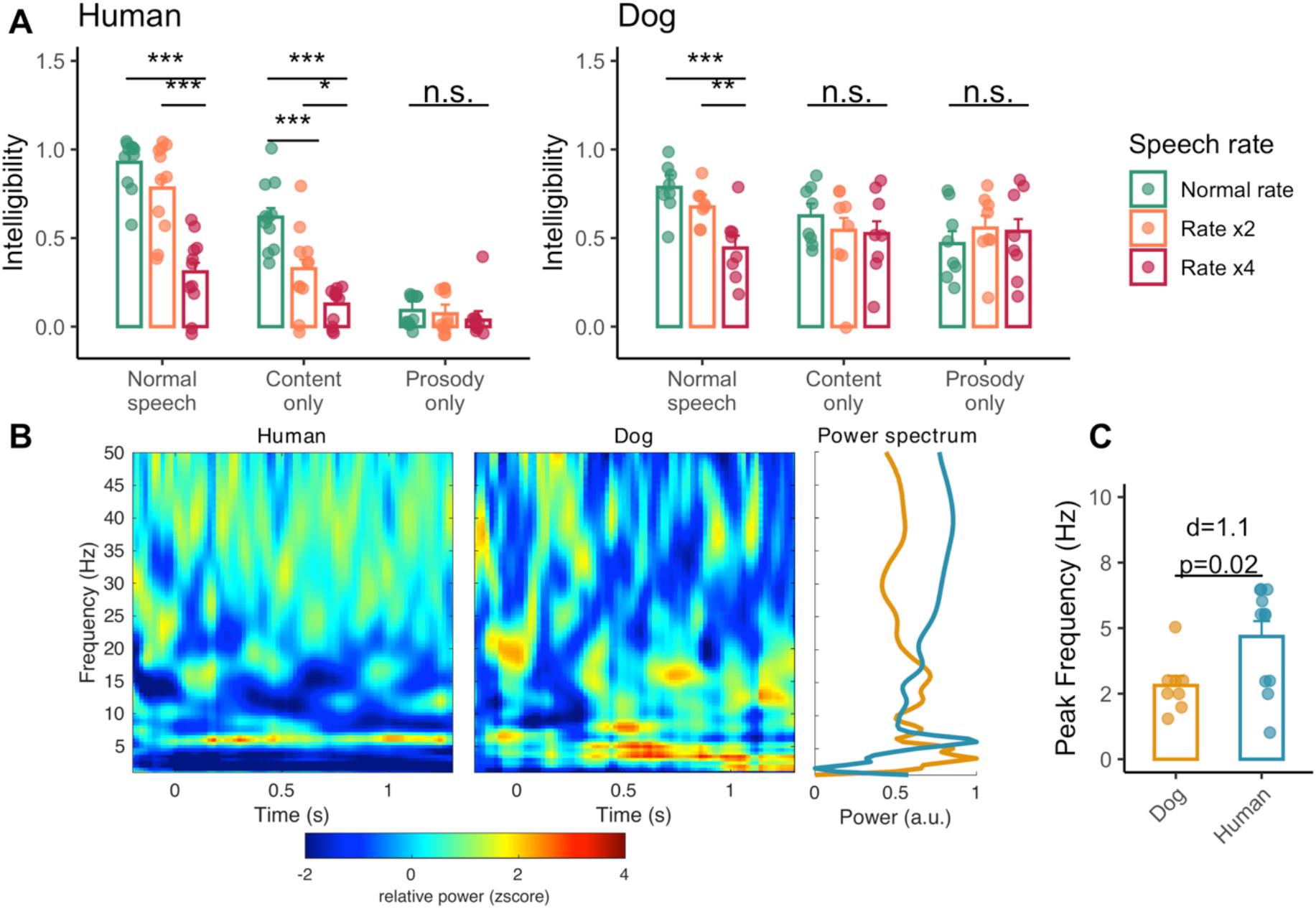
Speech stimulus alteration effects on intelligibility and characterisation of neural responses. A) Mean (± SE) behavioural responses according to speech type and rate in humans and dogs. Tukey-corrected, post-hoc pairwise comparisons are shown. *** p<0.001, ** p<0.01, * p<0.05. B) Time-frequency plots averaged across all conditions and individuals within species. Z-score transformed relative power is plotted to ease visual comparison across species. C) Power spectra (peak normalised and averaged between 0-1.3s) and unpaired t-test between species on frequency of highest power (range=1-7Hz).

We then quantified the two species’ neural responses, restricting the EEG analyses to the FCz electrode in humans and Cz in dogs, as they showed the strongest response to the acoustic stimuli (fig. S1, ^42,43^). Both dogs and humans showed increased power activity in the low frequency range (<10Hz, Figure 3B), confirming and characterising the auditory cortex activity reported in fMRI studies of dog speech processing ^44–46^. However, we noted a first difference between the two species’ neural responses in this frequency range. Dogs showed a predominant power increase in the delta band (1-3 Hz), as opposed to the theta band (4-7Hz) in humans (unpaired t-test: t=-2.47, df=17, p=0.02, Cohen’s d=1.1, Figure 3C), speaking to possible divergent auditory processes.

Given the presence of a stimulus-related and sustained neural response, we then established whether dogs display evidence of a speech tracking response under normal speech conditions. Cerebro-acoustic coherence, a measure that quantifies the phase-locking of neural signals to speech envelope ^18,19^, was above the mean random coherence value throughout the 1-10 Hz range in humans, but restricted to a 1-3 Hz peak in dogs (Figure 4A). Averaged coherence values in the delta band were significantly higher in the real pairings than the random ones in humans (paired t-test: t=-5, df=10, p<0.001, d=1.5) and in dogs (t=-3, df=7, p=0.02, d=1.1). However, theta coherence was significantly higher in humans (t=-2.87, df=10, p=0.02, d=0.9) but not in dogs (t=-0.6, df=7, p=0.5, d=0.2). In other words, dogs show evidence of auditory tracking capabilities, as do other species ^47–49^, however, in the context of speech stimulation, such tracking is restricted to the delta band (Figure 4B).

**Figure 4.**
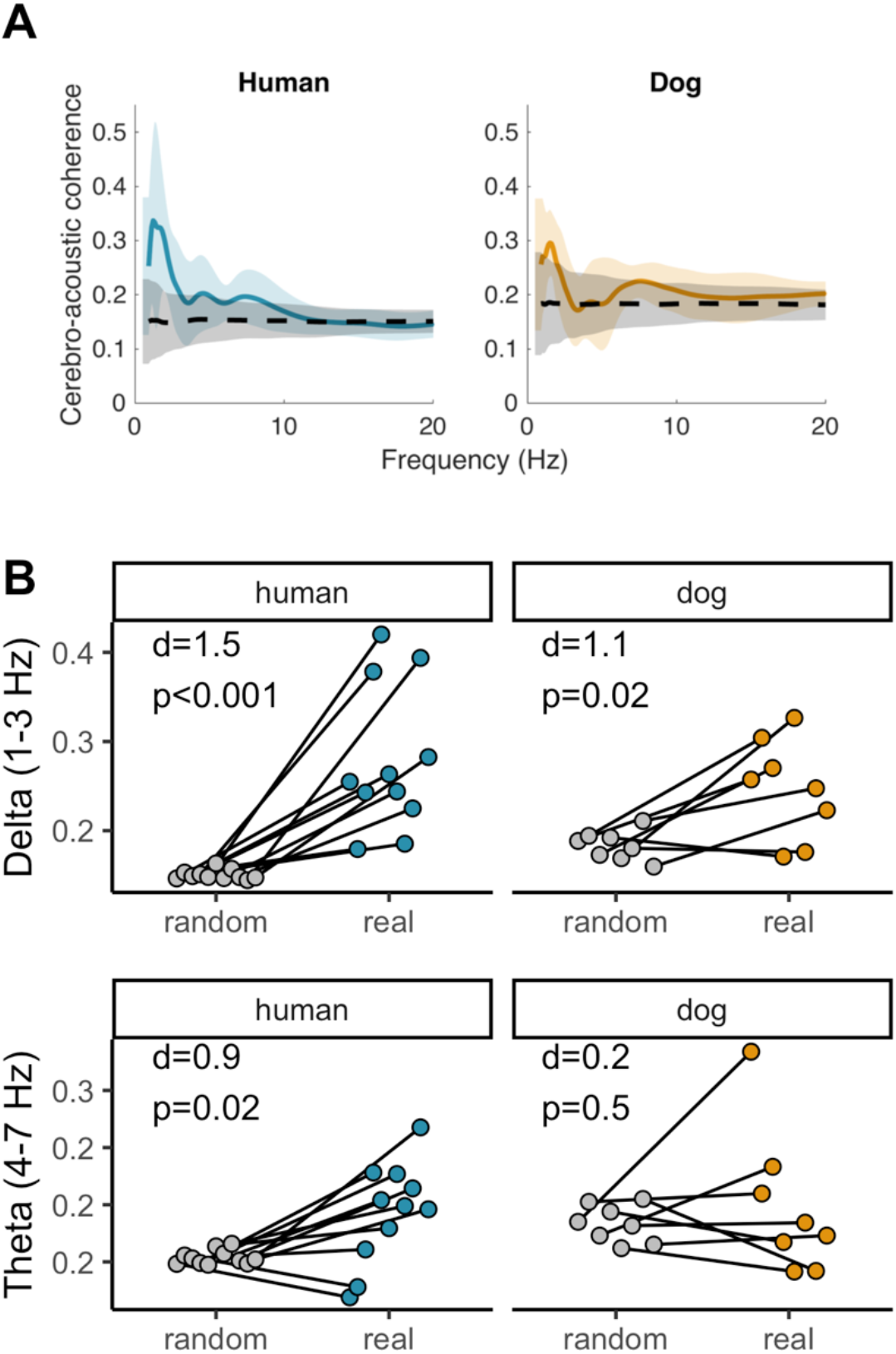
Speech tracking in both species and both the delta and the theta bands. A) Mean and SD cerebro-acoustic coherence over the 1-20 Hz range, calculated from the normal speech condition. Black dashed line shows mean and SD random coherence values for pairings of neural signals with randomised acoustic envelopes. B) Paired t-test of coherence in the delta and theta range between the real cerebro-acoustic and cerebro-randomised acoustic pairings.

Given that dogs and humans differ with regards to the primary frequency band that responds most strongly to speech stimuli (i.e. the delta band for dogs, and theta band for humans), we subsequently extracted the peak rhythm of each speech stimulus in these two bands, corresponding to the word and syllable rate, respectively, and the individual cerebro-acoustic coherence value for these specific rhythms (hereafter ‘word coherence’ and ‘syllable coherence’). We confirmed that increasing speech rate had a negative effect on cerebro-acoustic coherence in both species and at both granularity levels (fig. S2). In humans, both syllable and word coherence decreased as syllable rate (F_1,90.5_=9.2, p=0.003, fig. S2A) and word rate (F_1,84_=4.2, p=0.04, fig. S2B) increased respectively, but speech type had no effect in either model (Syllable model: F_2,83_=0.27, p=0.8; Word model: F_2,83_=0.5, p=0.6). The same pattern was found for dogs, with both syllable and word coherence dropping with increasing syllable rate (F_1,65.5_=5, p=0.03, fig. S2C) and word rate (F_1,61_=9.1, p=0.003, fig. S2D) respectively, while speech type (Syllable model: F_2,59_=1.5, p=0.2; Word model: F_2,59_=2.9, p=0.07) had no effect.

Remarkably however, the two species differed with regards to the granularity level at which tracking was most strongly related to behavioural outputs (Fig. 5A). Specifically, in humans, word coherence did not explain intelligibility (F_1,58.3_=1.3, p=0.3) while increased syllable coherence led to increased intelligibility (F_1,89.5_= 5.5, p=0.02). Conversely, in dogs syllable coherence had no impact on speech intelligibility (F_1,62_=0.21, p=0.6), while intelligibility increased with increasing word coherence (F_1,62_= 4.69, p=0.03). Interestingly, in both species the speech type main effect remained (humans syllable model: F_2,85_=49.9, p<0.001; dogs word model: F_2, 61_=3.2, p=0.05, Figure 5B), with significant differences among the intercepts of all speech types in humans (all pairwise comparisons: p<0.001) and higher intelligibility in the normal speech condition compared to the prosody-only condition (Norm. speech – Prosody-only est=0.1, df=69, t=2.4, p=0.05) in dogs (all other pairwise comparisons: p>0.05). In other words, like humans, dogs’ comprehension of speech appears to involve more than stimulus-driven auditory processes ^22,43,50^.

**Figure 5:**
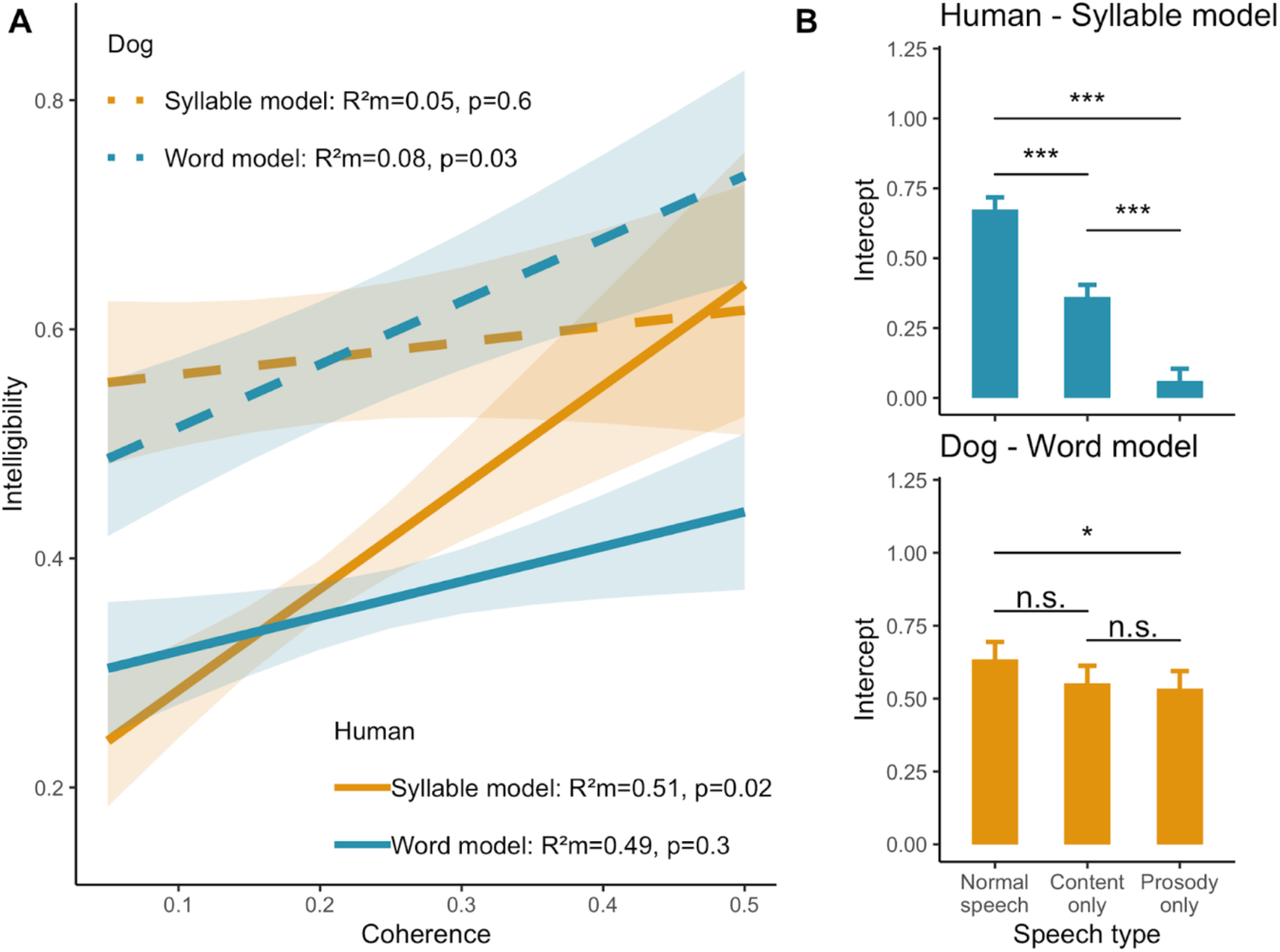
Increased speech neural tracking (cerebro-acoustic coherence) increases comprehension. A) Model slopes and 95% CI for the syllable and word coherence effect on intelligibility in dogs and humans. B) Mean and S.E. intercepts for each speech types in humans and dogs, showing that beyond speech tracking, additional processes must be present to explain the differences.

## DISCUSSION

While humans’ speaking rate corresponds to the frequency range of theta neural oscillations, dogs’ vocal rate aligns instead with the delta oscillation range. Further, dogs’ natural vocalisation rate is conserved across vocalisation types, not influenced by body weight and shows only limited inter-individual differences, suggesting that selective pressures favoured the maintenance of this regime. This is consistent with what is known from the speech production system, namely that despite wide linguistic variations, speech rate shows remarkable consistency cross-culturally ^11,51^. Thus, we propose that dogs, like humans, are subject to evolutionary factors that have kept production constrained within their species-specific ranges (Figure 1), possibly due to inherent differences in the function(s) of their respective communication systems ^4^. Interestingly, the theta vocal rhythm has been shown to be present in many primate species including both closely and more distantly-related ones ^52–54^ leading to hypotheses of an exaptation from masticatory movements ^55^. Yet, despite being masticators, dogs do not vocalize in that range and instead exhibit a lower rate, suggesting that the theta rhythm evolved sometime after the split between the Laurasiatheria and Euarchontoglires and begging for a more thorough characterization of the phylogeny of the theta rhythm and of the possible selective forces behind its emergence.

Importantly, we found three key pieces of evidence that lend support to the view that receiver sensory and neural systems are a primary selective force driving signal design ^22,57^. Dogs’ vocal rate is range-constrained in the delta band and during auditory perception of human speech, dogs primarily track slow amplitude modulations in speech that correspond to delta oscillations and only word-level predicts intelligibility (Figure 4 and 5). At this point, it should be noted that there is currently no data available on dogs’ processing of conspecific vocal sequences. Yet, it seems highly probable that dogs would similarly rely on delta tracking to process conspecifics signals given that cortical tracking is a basal processing mechanism ^47,48,56^ and that delta oscillations best match this species’ vocal temporal patterning as shown in this study. Furthermore, we found that rather than dogs syllabifying speech, humans modify their vocal output to approach dogs’ temporal channel. Thus, while the exact rhythm at which production systems align to reception systems is species-specific, such a process appears to also regulate adaptive heterospecific communication.

Further, whether when looking at the behavioural or neural level, we found that there existed differences relating to the type of speech broadcasted (Figure 3 and 5). Indeed, contrary to popular beliefs, we found no evidence that dogs primarily rely on ‘prosody’ rather than ‘content’ to respond to human vocal cues. Instead, successful responses require the fully integrated signal (Figure 3), confirming and extending previous results that used a preference-looking paradigm to investigate responses to ADS and DDS in dogs ^58^. To borrow a term from the multi-modal communication literature, while in humans, prosody and content have a modulatory effect, for dogs we see an emergence effect of the composite signal ^59^.

Finally, we found that speech tracking was not sufficient to fully explain intelligibility in both species, as differences in intercepts among speech types remained (Figure 5). In humans, speech comprehension relies on both bottom-up and top-down processes distributed across a wide range of cortical regions ^22,60,61^. Most notably, the former involves the hierarchical phase-amplitude coupling of theta and gamma frequency bands allowing for phoneme encoding ^15,62^ while the later involving motor cortex responses ^63^ that are causally related to perception ^64^. Arguably, both the need for and possibility to encode phonemes and the role of the motor cortex in predictive processes during auditory tasks likely mechanistically reflects within-species audio-motor tuning ^22^, and as such need not be speech specific ^65–67^. Yet, the dogs’ data also suggest that their processing of human speech entails more than just low-level auditory processing and while there is no data on dog neural coupling, a recent study mentioned differential activation in a cortical premotor region when comparing familiar vs unfamiliar language processing ^50^. Thus, it will be greatly interesting for future research to investigate whether and to what extend hierarchical bottom-up and predictive processes initially linked to within-species acoustic processing can adapt to or constrain inter-species communication.

Overall these results reveal that dogs’ auditory and vocal systems have aligned on a single temporal processing window that differs from that of humans, and which remains predominant even when dogs process and appropriately respond to human speech. In parallel, we show that humans who speak to their dogs adopt a speech rhythm that differs from adult-directed speech and more closely aligns with the dog’s neural delta oscillatory capacity. Thus, in the history of the dog-human relationship, it appears that the neural constraints of the dogs’ reception system have limited this heterospecific communication to a temporal structure falling midway between the natural speech rate and a slower rate that would perfectly match the dog’s analysis capacity.

## METHODS

### Ethics statement

All the dogs used in this study were pet dogs who lived with their caregivers. As the tests took place in France, involved non-invasive EEG recordings and behavioural tests, no ethical approval was required under the French law. The human participants all provided informed consent prior to the experiments and the procedures were approved by the ethics commission of Geneva University CUREG.202011.18.

### Subjects

#### Dogs

Dog owners were recruited by contacting canine clubs located in France. After initial contact with potential participants, dogs were recruited if they met the following inclusion criteria: being 1 year or older, having no hearing deficits, a good sociability level, high trainability and a good level of education. This recruitment process resulted in a pool of 12 dogs (seven females) aged 1-13 years old being included, all being medium to large dog breeds, the smallest being Shetland sheepdogs and the biggest being the Beauceron.

#### Humans

We also recruited the same number of human participants (six women) from the clubs who served as paired controls. Inclusion criteria for human participants were: being aged 18-65 years old, having no hearing deficit, no psychiatric or motor disorders and speaking French fluently.

### Procedure

#### Perception experiment

##### Pre-experiment dog training

We developed a training protocol using positive reinforcement and behavioural shaping, to condition dogs to wear the EEG equipment while remaining still. First, dogs were clicker-trained to lay down while resting their head. Once dogs could maintain this position for at least 15s, they were habituated to wear a headband (happy hoodie ®) normally used during toileting, to which we made holes to let the ears out (Figure 2, photo inserts). Then they were finally conditioned to maintain the position, while wearing the headband and listening to a variety of noises including music, environmental noises and voices. Dogs were judged sufficiently experienced once they could maintain this position regardless of noise or other environmental disturbances for at least 15s.

##### Acoustic stimuli

Typically, comprehension is assessed in humans by asking participants to rate word sequences on an intelligibility scale. As this was not possible for dogs, we selected words that the dogs had learned to respond to, i.e. command words (e.g. “sit”, “come”), allowing us to use behavioural responses to these words as an index of ‘comprehension’ in the sense of a successful stimulus-action relationship. For each dog, we recorded five command words spoken by their owners during a typical training session to obtain original, naturalistic DDS. The words were a mix of mono- and disyllabic words (Figure 2). Each dog listened (EEG task) and responded (behavioural task) to their specific set of command words. Their matched control human participant also listened to the same stimuli. Recordings were made with a Sennheiser ME64 microphone and a K6 module mounted onto a FOSTEX FR-2LE field recorder in 44.1kHz-16 bit wav format. One exemplar of each command word was selected based on the sound quality and on whether that occurrence resulted in a clear, successful behavioural response. The selected command words were first high-pass filtered at 100hz, independently normalised at −2dB and then concatenated into one word stream with 300 ± 50ms silent intervals in between command words. We then used PRAAT and the VocalToolkit plug-in to construct the acoustic stimuli. In total we constructed nine word stream stimuli using a fully crossed design of the three levels of speech type and the three levels of speech rate (Figure 2).

In the <u>Content-only</u> condition, we first changed the pitch median of the original dog-directed word sequence to match that of the owner’s adult-directed speech pitch, and to remove all pitch modulations. In a second step, we altered the intensity component of prosody, by reversing the natural intensity contour, while keeping the speech forward. To create the <u>Prosody-only</u> condition, we reversed each individual word rendering the speech unintelligible, while keeping their order in the word stream. Because this process also reversed pitch modulation and intensity, we then copied the pitch and intensity modulation patterns from the original speech word stream, effectively reinstating the original prosody. Finally, given that these procedures resulted in undesired contingent effects, such as slightly robotized voice effects, we also created a control <u>Normal speech</u> condition by first making the pitch monotone and recopying the original pitch contour from the original recording. This ensured that these contingent effects were also present in the Normal word stream and thus controlled for.

To accelerate the speech rate, we used the ‘change tempo’ function in Audacity®, https://audacityteam.org/, which accelerates the rate without impacting the pitch. Each stream was compressed by a factor of 2 (twice as fast) and a factor of 4 (four times as fast). This resulted in a total of nine word streams to which we affixed the dog’s name in its original form, as a way of capturing the dog’s attention throughout the experimental session. We then created experimental tracks that included all 9 word streams repeated 40 times each, presented in a random order and separated by an inter-stimulus silent interval of 1.5 ± 0.5s. Experimental tracks lasted on average 23 min.

##### Experimental location

All tests took place at the owner’s home whenever possible, or at another place familiar to the dog (such as another participant’s house belonging to the same canine club). This avoided having to familiarise animals with new locations and gave us more flexibility during the COVID-19 situation. Typically, this involved using the living room area of the house, with the dog being either positioned on a bedding or a couch, depending on its usual place during the pre-experiment training.

##### EEG listening task

On the day of the EEG test, dogs were fitted with 1 to 4 golden cup electrodes (at least Cz and if possible, C3, C4 and POz) using gel and a conductive paste, and connected to a g.Nautilus amplifier (g.tec ©) secured on the back of the dog, which wirelessly transmitted data to a receiver connected to a recording DELL laptop. The reference electrode was placed at the nap of the neck (Figure 2, photo inserts). Electrodes were then secured by the headband to prevent any movement during the experiment. Electrode impedance was kept under 30kΩ and data were recorded at a 500 Hz sampling rate. Dogs layed down facing a PREMIO 8 speaker (T.A.G Montpellier, France) placed 2m away. The experimental track was then broadcast at 60 ± 5dBC. The experiment was paused regularly to reward the dog for maintaining the position or when the dog became restless. On average the dog EEG listening task lasted 39.6 ± 15.8min.

Human recordings were made as similar as possible, using the same recording device and the same set-up. The only differences being that we used 7-8 gel-based g.SCARABEO (g.tec ©) active electrodes inserted in a cap and ear-referenced, and that the participants were asked to sit in a chair and instructed to avoid movements and blinking during the stimulus presentation. No breaks were given during the presentation.

### Behavioural task

#### Humans

Participants were asked before the EEG listening task, to score the linguistic material. For that, they were equipped with headphones and listened to each stimulus and were prompted to score on a scale of 0 to 5 how many words they understood. The word streams were randomly ordered but only presented once to avoid learning effects.

#### Dogs

To obtain a comparable index for dogs, we used a playback experiment where dogs were made to listen to each command word separately (45 words in total) and scored on how well they responded to the command. To do so, we installed the speaker at the mouth level of the dog owner, who stood quietly next to it while wearing sunglasses and a face mask, holding their arms along their body or behind their back. This procedure ensured that the experiment was as realistic as possible while preventing dogs from using visual cues to answer the command. Prior to each command word, the dog was positioned in front of the speaker 1-2 away in a position that allowed it to display the appropriate behavioural response (e.g. standing up if the next command was a ‘sit’). Each command word was played a maximum of three times with 10s of silent interval in between. The first time the word command was played, it was preceded by the name of the dog, to grab her attention and replicate typical training settings. After the command word was played, we scored on a scale of 1 to 5 how accurately the dog responded to the command (Table S2). If the dog obtained a score of 4 or 5 (i.e. perfect response within the 10s scoring interval), she was rewarded with her usual treat, the playback series for that command was stopped and we moved on to the next command word series, again mimicking a typical training session. Scoring was performed by the experimenter and the dog owner. If the two disagreed, the lower response score was given. If the dog became restless and/or inattentive, the experiment was interrupted by a play and/or walk session and then resumed. On average the task lasted 53.5 +/-20.3min.

### Production experiment

#### Dogs

We collected vocal sequences from youtube videos using the freely available Audio Set database ^68^. A total of 143 sequences (30 individuals) lasting >1.5s were extracted, spanning the range of basic vocalisations in canids (barks, growls, whines, snarls and howls ^38^). We categorised the dogs according to their body size as either small (a terrier-like dog or below) or large and to their age class (adult vs juvenile). Whenever available, we recorded the breed of the dog and obtained the corresponding mean breed weight using the American Kennel Club website (https://www.akc.org/). For the Cane corso, data were unavailable on the AKC website, so we used the French equivalent, the Societé Centrale Canine website (https://www.centrale-canine.fr). Finally, for those three individuals whose breed was known and who were pups, we first estimated the age (in months) of the pup from the video and then used the weight curve of the corresponding weight category provided in ^69^ to obtain the mean weight (50% centile) at that age.

#### Humans

To keep datasets as comparable as possible, we extracted ADS and DDS sequences from youtube videos (ADS: 106 sequences, 27 individuals, 10 women; DDS: 149 sequences, 22 individuals, 16 women). We selected speech sequences from five different languages: English, French, Italian, Japanese and Vietnamese to cover the range of stress-, syllable- and mora-timed speech patterns. DDS sentences included both typical praising and command sentences. For 12 individuals (nine women) that produced DDS sequences, we were able to match one DDS and one ADS exemplar (matched for duration), either extracted from the same video or by looking at other videos published by that user. For this analysis, we were not able to find matching ADS and DDS sequences in Vietnamese.

### Measurements

#### Perception experiment

##### Intelligibility index

For humans, the intelligibility score corresponded to the proportion of correctly comprehended words. To obtain a comparable index for dogs, we calculated the mean response score from the maximum behavioural score obtained in response to each command word of a given condition (Table S2), and then scaled this variable between 0 and 1.

##### Audio signal

We computed the speech envelope (from the onset of the first command word) using the Hilbert transform, low-pass filtered below 30Hz using an 8^th^ order butterworth filter, to extract, for each participant, the word and syllable rate for each of the nine word streams from the power spectrum of the envelope. These word and syllable rate variables were then z-scored and subsequently used as regressors in the statistical analyses investigating the relationships between neural, acoustic and behavioural data.

##### EEG data

All EEG pre-processing steps were done in MATLAB using the fieldtrip toolbox ^70^ and custom-written scripts. EEG data were bandpass filtered between 1 and 70 Hz and a DFT filter was applied at 50, 100 and 150 Hz. Signals were then epoched from 1s pre-stimulus to the end of the word sequence. Human data were re-referenced to average and an independent component analysis (ICA) was used to remove eye blink data. ICA was not used on dog data, as for most subjects (8 out of 12), we only had the Cz recording electrode. Artefact rejection (eye blinks, muscle and jumps) was automatically done using fieldtrip functions, with species-specific cut-off z-values (more stringent for humans). A final visual inspection of all trials was used to remove any other trial that failed the rejection procedure. During these initial procedures, we had to exclude three dogs and one human participant due to poor signal quality, leaving nine dogs and 11 humans.

### Electrode selection

For most dogs we could only analyse EEG data from Cz, as from the nine dogs remaining after the initial pre-processing steps, seven were equipped with only Cz. Thus, we decided to similarly restrict further data analyses to one electrode for humans. To select which electrode to keep we used a decoding approach, using the mTRF model ^71^. Briefly, mTRF models use regularised linear regression to find the latent relationships between the stimulus features (in our case the speech envelope) and the neural response. We ran mTRF models for each subject and each electrode separately, restricting the shifting lag from 100ms pre-stimulus onset to 500ms post-stimulus onset. We then calculated the correlation between the reconstructed and the actual stimulus and saved the mTRF r value obtained as our measure of how well each electrode responded to the task. A linear-mixed model with electrode as a fixed effect and subject ID as a random term, followed by post-hoc analyses showed that FCz had significantly higher correlation value compared to the other electrodes, and was thus selected for further analyses (fig. S1).

### Cerebro-acoustic coherence

To assess the extent of cortical phase-locking to the speech temporal structure, we used the cerebro-acoustic coherence index. Focusing on the control normal speech condition, we first obtained the cross-spectral density between neural signal and the speech envelope using a wavelet method between 1-20Hz in 0.1 Hz frequency steps and 0.01 ms time steps, from 0.6 s post stimulus onset to 1.3s. This time window was selected to exclude ERP components resulting from the first word of the sentence, which was always the dog’s name, and to allow keeping trial length equal across subjects. We then used the coherence function in fieldtrip to compute the phase coherence between the speech envelope and neural signal. To evaluate how well subjects tracked the speech signal we compared the actual coherence to random coherence values obtained from the pairings of neural data with randomised acoustic envelopes averaged over 100 runs. To further characterise neural tracking in the two most relevant auditory frequency bands, i.e. delta and theta bands, we extracted mean coherence values (delta: 1-3Hz; theta: 4-7Hz) in both real and random datasets and compared them using paired t-tests. Then, to explore how tracking was influenced by speech type and rate and how it related to behavioural data, we calculated, for each subject in each condition, the mean word and syllable coherence (time window: 0.6-1.3s post-stimulus onset, time steps: 0.01s, frequency steps: 0.5Hz) value centred around the subject-specific stimulus word and syllable rate (+/-0.5Hz).

### Production experiment

#### Vocal rate and dominant acoustic frequency

Acoustic analyses were performed using the seewave package in R (Sueur et al. 2008). To extract the peak vocal rate, i.e. the predominant rhythm at which vocalisations in a sequence are produced, we first bandpass filtered the sequence between 0.1-10kHz and then computed the signal’s envelope using the Hilbert transform. This envelope was further low-pass filtered below 20Hz using a 4th order butterworth filter and a wavelet method was used to obtain the frequency decomposition of the signal and extract the frequency of the highest peak.

As a control analysis, we also extracted the dominant acoustic frequency of one vocalisation per sequence (selected based on its signal-to-noise ratio) for the dogs, and the sentence’s mean fundamental frequency (F0) for humans. For the dogs, the vocal unit was first bandpass filtered between 50Hz-2kHz, and the averaged frequency spectrum was then computed to extract the frequency of the peak amplitude. We focused on the dominant acoustic frequency rather than the fundamental frequency, because the latter is not always quantifiable, particularly in noisy and chaotic vocalisations such as barks. For humans, we used PRAAT (with standard settings) to extract the mean F0 in each sequence, as in human speech, F0 is both easy to compute and a better characterisation of pitch than dominant frequency.

#### Potential for individual coding (PIC)

To assess potential individual-related variations in the acoustic parameters, we calculated the within- and between-individual coefficients of variation (CVw and CVb respectively) using the formula for small samples (Sokal and Rohlf 1995) and obtained the feature’s PIC value, which is the CVb/meanCVw ratio, where meanCVw is the mean value of the CVw for all individuals ^72^.

### Statistical analyses

#### Behavioural and EEG data

To investigate how the experimental conditions influenced neural and behavioural responses, we used Linear-Mixed Models (LMMs). Models always included participant ID as a random term. Fixed effects varied depending on the question being addressed and were always first specified as the full model, then interaction terms were dropped if they did not reach significance. Statistical significance of fixed effects was assessed using F tests and the Kenward-Roger method of degrees-of-freedom approximation, as it has been shown to be a reliable method when LMMs are balanced ^73^. Post-hoc pairwise comparisons were Tukey corrected. Visual inspection of plots showed that the normality and homoscedasticity of residuals and random effects assumptions were met in all cases. For full reporting of these models and all other statistical analyses, refer to the sup. mat. Rmd document. Unpaired or paired t-tests were used in comparisons when they were warranted.

#### Vocal production data

We tested for differences in vocal rates between dogs and humans, using a LMM with species as a fixed effect and subjects within call/language types as random effects. We used LMMs to assess whether the acoustic measurements varied with call/language type adding weight class and subject ID as random terms in dogs, while in humans, the random effects were subject ID and sex. Finally, to assess whether acoustic parameters were allometrically related to body weight in dogs, we first log-transformed the variables and then used an LMM with vocal class and subject ID as random terms. Unpaired or paired t-tests were used in comparisons where they were warranted.

## DATA AVAILABILITY

Data and codes are available here: figshare DOI (DOI number will be made available at the time of publication)

## ACKNOWLEDGEMENTS

We are thankful to Silvia Marchesotti and Johanna Nicolle for their help in the initial piloting stage of this work. This research received funding from the NCCR Evolving Language, Swiss National Science Foundation Agreement #51NF40_180888.

## AUTHORS CONTRIBUTIONS

*Experimental design*: ED, FG & ALG, *Experimental setup:* ED & TL*, Data collection and experiments:* ED & TP, *Data analysis:* EC, LA, TP & ALG, *Manuscript preparation:* ED, LA & ALG, *Manuscript revision: all co-authors*.

## MATERIALS & CORRESPONDANCE

All requests should be sent to E. Déaux.

## References

1 Charlton, B. D. & Reby, D. The evolution of acoustic size exaggeration in terrestrial mammals. Nature communications 7, 12739 (2016).

2 Taylor, A. M. & Reby, D. The contribution of source–filter theory to mammal vocal communication research. Journal of Zoology 280, 221–236, doi:10.1111/j.1469-7998.2009.00661.x (2010).

3 Ryan, M. J., Fox, J. H., Wilczynski, W. & Rand, A. S. Sexual selection for sensory exploitation in the frog *Physalaemus pustulosus*. Nature 343, 66–67, doi:10.1038/343066a0 (1990).

4 Arnal, L. H., Flinker, A., Kleinschmidt, A., Giraud, A. L. & Poeppel, D. Human screams occupy a privileged niche in the communication soundscape. Curr Biol 25, 2051–2056, doi:10.1016/j.cub.2015.06.043 (2015).

5 Hoy, R. R., Pollack, G. S. & Moiseff, A. Species-Recognition in the Field Cricket, *Teleogryllus oceanicus*: Behavioral and Neural Mechanisms. American Zoologist 22, 597–607, doi:10.1093/icb/22.3.597 (2015).

6 Ghazanfar, A. A., Smith-Rohrberg, D. & Hauser, M. D. The role of temporal cues in rhesus monkey vocal recognition: Orienting asymmetries to reversed calls. Brain, Behavior and Evolution 58, 163–172, doi:10.1159/000047270 (2001).

7 Blumstein, D. T. & Armitage, K. B. Alarm calling in yellow-bellied marmots: I. The meaning of situationally variable alarm calls. Animal Behaviour 53, 143–171 (1997).

8 Drăgănoiu, T. I., Nagle, L. & Kreutzer, M. Directional female preference for an exaggerated male trait in canary (*Serinus canaria*) song. Proceedings of the Royal Society of London. Series B: Biological Sciences 269, 2525–2531, doi:10.1098/rspb.2002.2192 (2002).

9 Galeotti, P., Sacchi, R., Rosa, D. P. & Fasola, M. Female preference for fast-rate, high-pitched calls in Hermann’s tortoises *Testudo hermanni*. Behavioral Ecology 16, 301–308, doi:10.1093/beheco/arh165 (2005).

10 Pauly, G. B., Bernal, X. E., Rand, A. S. & Ryan, M. J. The vocal sac increases call rate in the Túngara frog *Physalaemus pustulosus*. Physiological and Biochemical Zoology 79, 708–719, doi:10.1086/504613 (2006).

11 Coupé, C., Oh, Y. M., Dediu, D. & Pellegrino, F. Different languages, similar encoding efficiency: Comparable information rates across the human communicative niche. Science Advances 5, eaaw2594, doi:10.1126/sciadv.aaw2594 (2019).

12 Poeppel, D. & Assaneo, M. F. Speech rhythms and their neural foundations. Nature Reviews Neuroscience 21, 322–334, doi:10.1038/s41583-020-0304-4 (2020).

13 MacNeilage, P. F., Davis, B. L., Kinney, A. & Matyear, C. L. The motor core of speech: A comparison of serial organization patterns in infants and languages. Child development 71, 153–163 (2000).

14 Schroeder, C. E., Wilson, D. A., Radman, T., Scharfman, H. & Lakatos, P. Dynamics of Active Sensing and perceptual selection. Current Opinion in Neurobiology 20, 172–176, 10.1016/j.conb.2010.02.010 (2010).

15 Hyafil, A., Fontolan, L., Kabdebon, C., Gutkin, B. & Giraud, A.-L. Speech encoding by coupled cortical theta and gamma oscillations. Elife 4, e06213 (2015).

16 Ahissar, E. et al. Speech comprehension is correlated with temporal response patterns recorded from auditory cortex. Proceedings of the National Academy of Sciences 98, 13367–13372 (2001).

17 Luo, H. & Poeppel, D. Phase patterns of neuronal responses reliably discriminate speech in human auditory cortex. Neuron 54, 1001–1010, doi:10.1016/j.neuron.2007.06.004 (2007).

18 Peelle, J. E., Gross, J. & Davis, M. H. Phase-locked responses to speech in human auditory cortex are enhanced during comprehension. Cereb Cortex 23, 1378–1387, doi:10.1093/cercor/bhs118 (2013).

19 Doelling, K. B., Arnal, L. H., Ghitza, O. & Poeppel, D. Acoustic landmarks drive delta-theta oscillations to enable speech comprehension by facilitating perceptual parsing. Neuroimage 85 Pt 2, 761–768, doi:10.1016/j.neuroimage.2013.06.035 (2014).

20 Pefkou, M., Arnal, L. H., Fontolan, L. & Giraud, A. L. theta-band and beta-band neural activity reflects independent syllable tracking and comprehension of time-compressed speech. J Neurosci 37, 7930–7938, doi:10.1523/JNEUROSCI.2882-16.2017 (2017).

21 Doelling, K. B., Arnal, L. H. & Assaneo, M. F. Adaptive oscillators provide a hard-coded Bayesian mechanism for rhythmic inference. bioRxiv 2022.2006.2018.496664, doi:10.1101/2022.06.18.496664 (2022).

22 Giraud, A. L. & Poeppel, D. Cortical oscillations and speech processing: emerging computational principles and operations. Nat Neurosci 15, 511–517, doi:10.1038/nn.3063 (2012).

23 Hare, B. & Tomasello, M. Human-like social skills in dogs? Trends in Cognitive Sciences 9, 439–444, 10.1016/j.tics.2005.07.003 (2005).

24 Andics, A., Gacsi, M., Farago, T., Kis, A. & Miklosi, A. Voice-sensitive regions in the dog and human brain are revealed by comparative fMRI. Curr Biol 24, 574–578, doi:10.1016/j.cub.2014.01.058 (2014).

25 Pilley, J. W. & Reid, A. K. Border collie comprehends object names as verbal referents. Behav Processes 86, 184–195, doi:10.1016/j.beproc.2010.11.007 (2011).

26 Kaminski, J., Call, J. & Fischer, J. Word learning in a domestic dog: evidence for “fast mapping”. Science 304, 1682–1683 (2004).

27 Boros, M. et al. Neural processes underlying statistical learning for speech segmentation in dogs. Current Biology 31, 5512–5521.e5515, 10.1016/j.cub.2021.10.017 (2021).

28 Ratcliffe, V. F. & Reby, D. Orienting asymmetries in dogs’ responses to different communicatory components of human speech. Curr Biol 24, 2908–2912, doi:10.1016/j.cub.2014.10.030 (2014).

29 Fukuzawa, M., Mills, D. S. & Cooper, J. J. The effect of human command phonetic characteristics on auditory cognition in dogs (Canis familiaris). J Comp Psychol 119, 117–120, doi:10.1037/0735-7036.119.1.117 (2005).

30 Fitch, T. The phonetic potential of nonhuman vocal tracts: Comparative cineradiographic observations of vocalizing animals. Phonetica 57, 205–218, doi:10.1159/000028474 (2000).

31 Lieberman, P. Vocal tract anatomy and the neural bases of talking. Journal of Phonetics 40, 608–622 (2012).

32 Boë, L.-J. et al. Which way to the dawn of speech?: Reanalyzing half a century of debates and data in light of speech science. Science Advances 5, eaaw3916 (2019).

33 Burnham, D., Kitamura, C. & Vollmer-Conna, U. What’s new, pussycat? On talking to babies and animals. Science 296, 1435–1435, doi:10.1126/science.1069587 (2002).

34 Hirsh-Pasek, K. & Treiman, R. Doggerel: Motherese in a new context. Journal of Child Language 9, 229–237, doi:10.1017/S0305000900003731 (1982).

35 Ben-Aderet, T., Gallego-Abenza, M., Reby, D. & Mathevon, N. Dog-directed speech: why do we use it and do dogs pay attention to it? Proceedings of the Royal Society B: Biological Sciences 284, 20162429, doi:10.1098/rspb.2016.2429 (2017).

36 Leong, V., Kalashnikova, M., Burnham, D. & Goswami, U. The temporal modulation structure of infant-directed speech. Open Mind 1, 78–90, doi:10.1162/OPMI_a_00008 (2017).

37 Kalashnikova, M., Peter, V., Di Liberto, G. M., Lalor, E. C. & Burnham, D. Infant-directed speech facilitates seven-month-old infants’ cortical tracking of speech. Scientific Reports 8, 13745, doi:10.1038/s41598-018-32150-6 (2018).

38 Cohen, J. & Fox, M. Vocalizations in wild canids and possible effects of domestication. Behavioural Processes 1, 77–92 (1976).

39 Yin, S. & McCowan, B. Barking in domestic dogs: context specificity and individual identification. Animal behaviour 68, 343–355 (2004).

40 Molnár, C., Pongrácz, P., Faragó, T., Dóka, A. & Miklósi, Á. Dogs discriminate between barks: The effect of context and identity of the caller. Behavioural Processes 82, 198–201, doi:10.1016/j.beproc.2009.06.011 (2009).

41 Bowling, D. et al. Body size and vocalization in primates and carnivores. Scientific reports 7, 41070 (2017).

42 Howell, T. J., Conduit, R., Toukhsati, S. & Bennett, P. Auditory stimulus discrimination recorded in dogs, as indicated by mismatch negativity (MMN). Behav Processes 89, 8–13, doi:10.1016/j.beproc.2011.09.009 (2012).

43 Magyari, L., Huszár, Z., Turzó, A. & Andics, A. Event-related potentials reveal limited readiness to access phonetic details during word processing in dogs. Royal Society Open Science 7, 200851, doi:10.1098/rsos.200851 (2020).

44 Andics, A. et al. Neural mechanisms for lexical processing in dogs. Science 353, 1030–1032 (2016).

45 Gábor, A. et al. Multilevel fMRI adaptation for spoken word processing in the awake dog brain. Scientific Reports 10, 11968, doi:10.1038/s41598-020-68821-6 (2020).

46 Bálint, A., Szabó, Á., Andics, A. & Gácsi, M. Dog and human neural sensitivity to voicelikeness: A comparative fMRI study. NeuroImage 265, 119791 (2023).

47 Lakatos, P., Gross, J. & Thut, G. A new unifying account of the roles of neuronal entrainment. Current Biology 29, R890–R905, 10.1016/j.cub.2019.07.075 (2019).

48 Boari, S., Mindlin, G. B. & Amador, A. Neural oscillations are locked to birdsong rhythms in canaries. European Journal of Neuroscience 55, 549–565, 10.1111/ejn.15552 (2022).

49 Theunissen, F. E. & Shaevitz, S. S. Auditory processing of vocal sounds in birds. Current opinion in neurobiology 16, 400–407 (2006).

50 Cuaya, L. V., Hernández-Pérez, R., Boros, M., Deme, A. & Andics, A. Speech naturalness detection and language representation in the dog brain. NeuroImage 248, 118811, 10.1016/j.neuroimage.2021.118811 (2022).

51 Pellegrino, F., Coupé, C. & Marsico, E. A cross-language perspective on speech information rate. Language, 539–558 (2011).

52 Morrill, R. J., Paukner, A., Ferrari, P. F. & Ghazanfar, A. A. Monkey lipsmacking develops like the human speech rhythm. Developmental Science 15, 557–568, 10.1111/j.1467-7687.2012.01149.x (2012).

53 Risueno-Segovia, C. & Hage, S. R. Theta Synchronization of Phonatory and Articulatory Systems in Marmoset Monkey Vocal Production. Current Biology, 10.1016/j.cub.2020.08.019 (2020).

54 Pereira, A. S., Kavanagh, E., Hobaiter, C., Slocombe, K. E. & Lameira, A. R. Chimpanzee lip-smacks confirm primate continuity for speech-rhythm evolution. Biology Letters 16, 20200232 (2020).

55 MacNeilage, P. F. The frame/content theory of evolution of speech production. Behavioral and Brain Sciences 21, 499–511, doi:10.1017/S0140525X98001265 (1998).

56 Doelling, K. B. & Poeppel, D. Cortical entrainment to music and its modulation by expertise. Proc Natl Acad Sci U S A 112, E6233–6242, doi:10.1073/pnas.1508431112 (2015).

57 Ghitza, O. The theta-syllable: a unit of speech information defined by cortical function. Frontiers in Psychology 4, doi:10.3389/fpsyg.2013.00138 (2013).

58 Benjamin, A. & Slocombe, K. ‘Who’s a good boy?!’ Dogs prefer naturalistic dog-directed speech. Animal Cognition 21, 353–364, doi:10.1007/s10071-018-1172-4 (2018).

59 Partan, S. R. & Marler, P. Issues in the classification of multimodal communication signals. The American Naturalist 166, 231–245 (2005).

60 Ghitza, O. Linking Speech Perception and Neurophysiology: Speech Decoding Guided by Cascaded Oscillators Locked to the Input Rhythm. Frontiers in Psychology 2, doi:10.3389/fpsyg.2011.00130 (2011).

61 Hovsepyan, S., Olasagasti, I. & Giraud, A.-L. Combining predictive coding and neural oscillations enables online syllable recognition in natural speech. Nature Communications 11, 3117, doi:10.1038/s41467-020-16956-5 (2020).

62 Lizarazu, M., Carreiras, M. & Molinaro, N. Theta-gamma phase-amplitude coupling in auditory cortex is modulated by language proficiency. Human Brain Mapping 44, 2862–2872, 10.1002/hbm.26250 (2023).

63 Cheung, C., Hamilton, L. S., Johnson, K. & Chang, E. F. The auditory representation of speech sounds in human motor cortex. elife 5, e12577 (2016).

64 Meister, I. G., Wilson, S. M., Deblieck, C., Wu, A. D. & Iacoboni, M. The Essential Role of Premotor Cortex in Speech Perception. Current Biology 17, 1692–1696, 10.1016/j.cub.2007.08.064 (2007).

65 Williams, H. & Nottebohm, F. Auditory responses in avian vocal motor neurons: a motor theory for song perception in birds. Science 229, 279–282 (1985).

66 Archakov, D. et al. Auditory representation of learned sound sequences in motor regions of the macaque brain. Proceedings of the National Academy of Sciences 117, 15242–15252, doi:10.1073/pnas.1915610117 (2020).

67 Kikuchi, Y. et al. Sequence learning modulates neural responses and oscillatory coupling in human and monkey auditory cortex. PLOS Biology 15, e2000219, doi:10.1371/journal.pbio.2000219 (2017).

68 Gemmeke, J. F. et al. in 2017 IEEE international conference on acoustics, speech and signal processing (ICASSP). 776–780 (IEEE).

69 Salt, C. et al. Growth standard charts for monitoring bodyweight in dogs of different sizes. PLOS ONE 12, e0182064, doi:10.1371/journal.pone.0182064 (2017).

70 Oostenveld, R., Fries, P., Maris, E. & Schoffelen, J.-M. FieldTrip: open source software for advanced analysis of MEG, EEG, and invasive electrophysiological data. Computational intelligence and neuroscience 2011, 1–9 (2011).

71 Crosse, M. J., Di Liberto, G. M., Bednar, A. & Lalor, E. C. The Multivariate Temporal Response Function (mTRF) Toolbox: A MATLAB Toolbox for Relating Neural Signals to Continuous Stimuli. Frontiers in Human Neuroscience 10, doi:10.3389/fnhum.2016.00604 (2016).

72 Robisson, P., Aubin, T. & Bremond, J.-C. Individuality in the voice of the emperor penguin *Aptenodytes forsteri*: adaptation to a noisy environment. Ethology 94, 279–290 (1993).

73 Schaalje, G. B., McBride, J. B. & Fellingham, G. W. Adequacy of approximations to distributions of test statistics in complex mixed linear models. Journal of Agricultural, Biological, and Environmental Statistics 7, 512–524, doi:10.1198/108571102726 (2002).

