## supplementary files for "Dogs’ sensory-motor tuning shapes dog-human vocal interactions"

**SUPPLEMENTARY FIGURES**

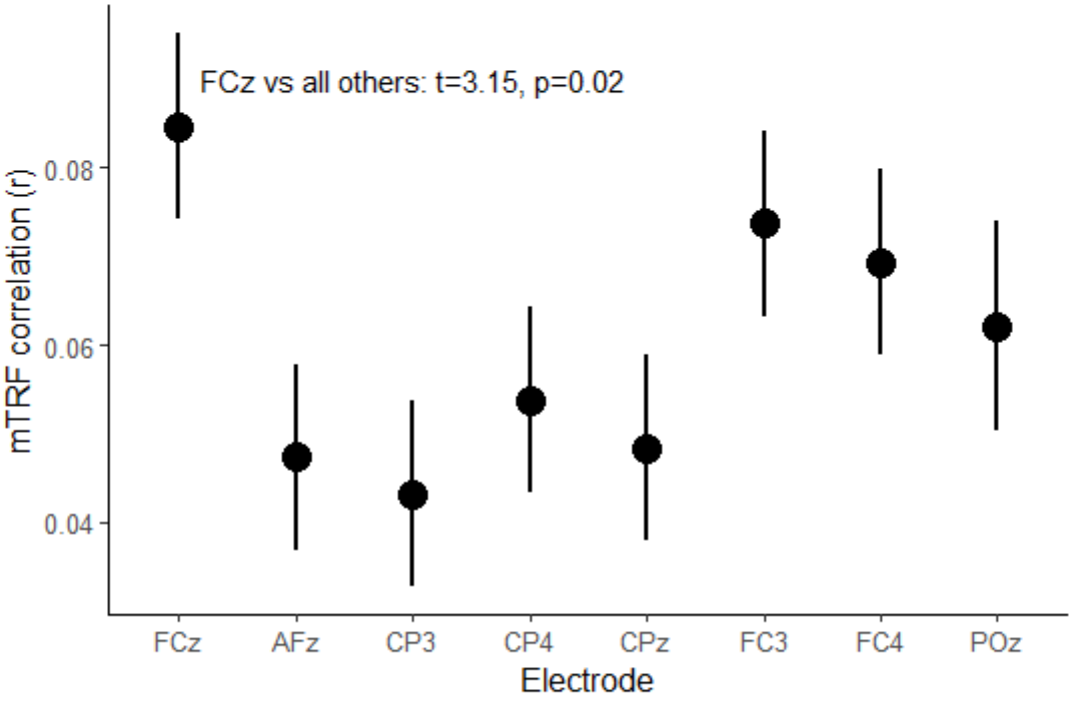


**Figure S1. Mean + SE mTRF r values for each electrode, from the human neural data**. The mTRF r values were obtained by first using a decoding model to reconstruct acoustic stimuli from neural data then by correlating this reconstructed acoustic data to the actual stimulus envelope. Thus, higher r values indicate that reconstructed data from that electrode better match the original stimulus. A linear-mixed model using electrodes as fixed effects and human ID as a random term, revealed significant differences among electrodes ($F_{7,67.1}$=3.17, p=0.006). Post-hoc tests (FDR corrected) were done to compare the mean value of one electrode to the average value of all other electrodes. FCz was the only electrode that showed significantly higher mTRF r values compared to all others.

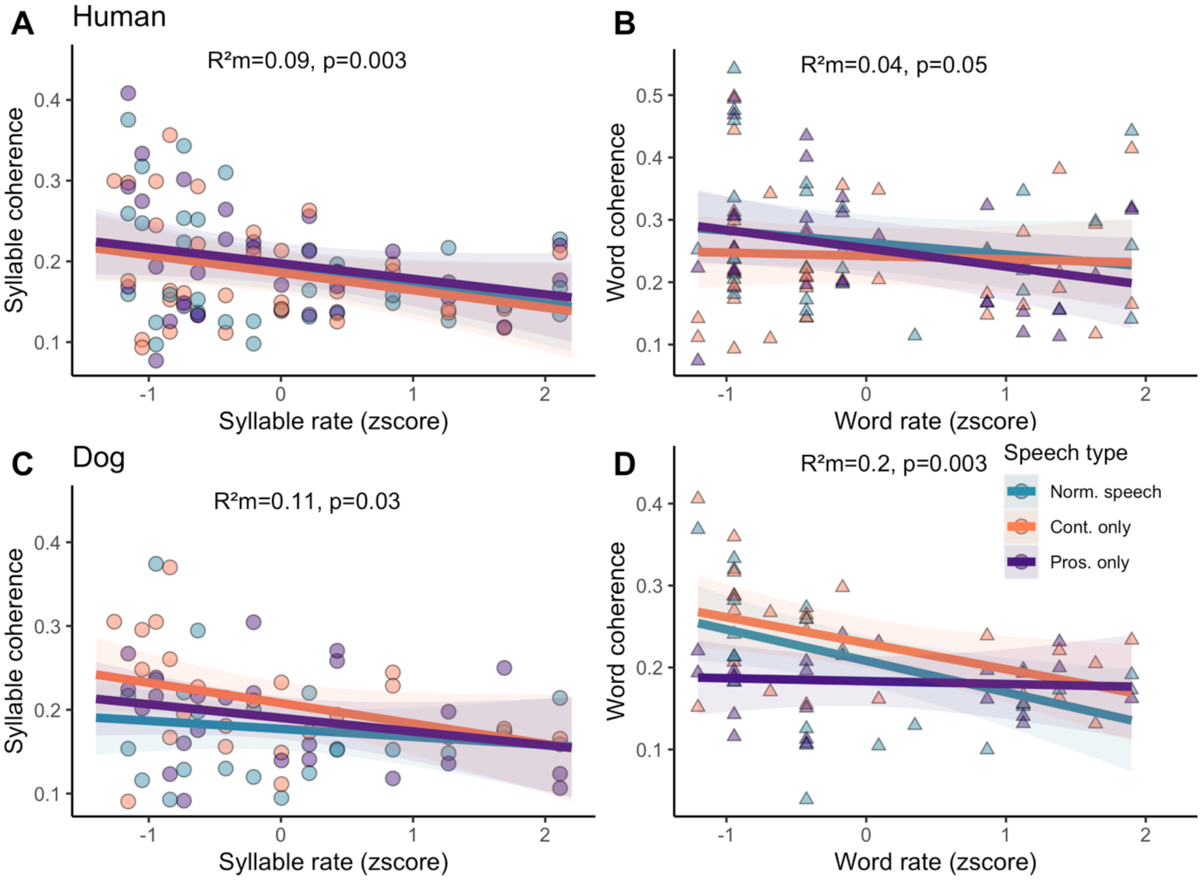


**Fig S2. Effect of speech rate on speech neural tracking (cerebro-acoustic coherence).** A) Slope estimate and 95% CI of syllable rate effect on syllabic coherence in humans for each speech type. B) Slope estimate and 95% CI of word rate effect on word coherence in humans for each speech type. C) Slope estimate and 95% CI of syllable rate effect on syllabic coherence in dogs for each speech type. D) Slope estimate and 95% CI of word rate effect on word coherence in dogs for each speech type.

**SUPPLEMENTARY TABLES**

**Table S1. Summary statistics of vocal rate (VR) and dominant acoustic frequency (DF) in dog vocalisations.** The potential for individual coding (PIC) is a measure that quantifies the ratio between the inter- and the intra-individual coefficients of variation, with values >1 indicating high individual distinctiveness. Typical vocal contexts are provided, although in some cases (e.g. barks, howls) vocalisations can be used in a range of situations spanning the affiliative-agonistic continuum.

| Vocal Class | Context | Mean (SD) VR (Hz) | Mean (SD) DF (Hz) | PIC VR | PIC DF | N |
|---|---|---|---|---|---|---|
| Bark | Alarm | 2.07 (0.68) | 663 (228) | 1.32 | 2.14 | 54 |
| Growl | Agonistic | 1.62 (0.78) | 502 (288) | 1.07 | 1.48 | 18 |
| Howl | Ambivalent | 1.83 (1.16) | 459 (163) | 1.1 | 1.6 | 33 |
| Snarl | Agonistic | 2.97 (1.55) | 795 (476) | 0.98 | 1.39 | 17 |
| Whine | Affiliative | 1.96 (1.3) | 857 (272) | 1.94 | 1.2 | 21 |

**Table S2.** Scoring scale for dogs' behavioural responses to command words.

| **Score** | **Behavioural response** |
|---|---|
| **5** | Complete response within 5s |
| **4** | Complete response after 5s and before 10s |
| **3** | Incomplete response |
| **2** | Non-specific or wrong response |
| **1** | No response within 10s |
